# Host-parasite interactions in perpetual darkness: macroparasite diversity in the cavefish *Astyanax mexicanus*

**DOI:** 10.1101/2023.05.16.540976

**Authors:** Ana Santacruz, David Hernández-Mena, Ramses Miranda-Gamboa, Gerardo Pérez-Ponce De León, Claudia Patricia Ornelas-García

## Abstract

*Astyanax mexicanus* has repeatedly colonized cave environments, displaying evolutionary parallelisms in many troglobitic traits. Despite being a model system for the study of adaptation to life in perpetual darkness, parasites infecting cavefish are practically unknown. In this study, we investigated the macroparasite communities of 18 cavefish populations from independent lineages and compared them with the parasite diversity of their sister surface fish populations, with the aim of better understanding the role that parasites play in the colonization of new environments. Thirteen parasite taxa were found in cavefish populations, including a subset of 10 of the 27 parasite taxa known for the surface populations. Parasites infecting the cavefish belong to five taxonomic groups: trematodes, monogeneans, nematodes, copepods, and acari. Monogeneans are the most dominant group, found in 14 caves. Macroparasites include species with direct life cycles and some trophically-transmitted parasites, including invasive species. Surprisingly, cave vs surface paired comparisons indicate higher parasite richness in the caves. The spatial variation in parasite composition across the caves suggests historical and geographical contingencies of the host-parasite colonization and the potential evolution of local adaptations. Base-line data on parasite diversity of cavefish populations of *A. mexicanus* sets the ground to explore the role of divergent parasite infections under contrasting ecological pressures (cave vs. surface environments) in the evolution of cave adaptive traits.

## INTRODUCTION

Host and parasites are engaged in complex interactions of constant reciprocal adaptation, imposing strong selective forces on each other, in some cases influencing their evolutionary trajectories (e.g. Bush et al., 2019). During the colonization of a new environment, hosts may lose parasites (enemy release hypothesis; Colautti et al., 2004) and maintain a subset of their original diversity, generating new parasite assemblages (Hoberg et al., 2012). Changes in these dynamics can alter the host-parasite interactions (Wolinska et al., 2008; Best et al., 2017), and produce rapid adaptations (Eizaguirre et al., 2012). Parasite selective pressures have implications for natural and sexual selection, for example, behavior modulation could change host habitat selection to avoid or promote parasite infection (Eizaguirre & Lenz, 2010; Mikheev et al., 2013; Demandt et al., 2018; Jolles et al., 2020), or parasite-influenced assortative mating could directly or indirectly influence mate choice (Milinski, 2014).

Differences in parasite infections are commonly associated with trophic disparity or environmental filters leading to local adaptations (Wegner et al., 2003; Karvonen & Seehausen, 2012; Roose et al., 2017). For instance, hosts may display defensive mechanisms to resist unique parasitic infections in the ecosystem (Eizaguirre et al., 2011, 2012), which could result in positive or negative fitness effects on the residents and/or immigrants and hybrids (Karvonen & Seehausen, 2012), even limiting the genetic flow between populations (Erin et al., 2019). For parasites, the complexity of its transmission strategy (i.e. direct or indirect life cycles), and host life history, using only aquatic hosts (autogenic life cycle) or cycling through aquatic and terrestrial hosts (allogenic life cycle), are main factors for dispersal and successful colonization (Criscione & Blouin, 2004). Trophically-transmitted species (indirect life cycle) rely heavily on suitable hosts to complete their life-cycle. However, in ecosystems with low nutrient availability, the opportunities for host-shifts are reduced. Instead, for parasites with direct life cycles, the host itself represents the environment (Lymbery, 2015), acting as a buffer for the challenging external conditions. Therefore, the trade-off with the environment can shape the evolution of the host-parasite interaction (Brunner & Eizaguirre, 2016; Brunner et al., 2017).

*Astyanax mexicanus* displays an extraordinary evolvability, allowing the repeated colonization of cave environments, characterized by low food availability and perpetual darkness. Across different subterranean rivers, *A. mexicanus* has evolved parallel phenotypes typical of troglobitic organisms: lack of eyes and pigmentation. Currently, 35 cave populations are known, which are located in Northeastern Mexico along three mountain ranges (Espinasa et al., 2018, 2020; Elliot et al., 2019, Miranda-Gamboa et al., 2023), acting as biogeographic barriers for the fish divergence and delimiting their phylogeographic patterns (Herman et al., 2018; Garduño Sánchez et al., 2022). The two independent lineages of cave and surface populations correspond to independent waves of colonization in a recent temporal scale (Strecker et al., 2004; Ornelas-García et al., 2008; Herman et al., 2018), which are referred as ‘Lineage 1’ and ‘Lineage 2’ (Garduño et al., 2022; Moran et al., 2023). One of the many highlights that have made *A. mexicanus* a fascinating model is that cavefish are still interfertile with the surface-dwelling fish, maintaining natural hybrid populations (Moran et al., 2022), and allowing studies about the inheritance of cave adaptive traits. Cavefish adaptations to perpetual darkness include morphological, behavioral, and physiological modifications (Wilkens & Strecker, 2017; Hyacinthe et al., 2019; Gross & Powers, 2020; Jeffery, 2020; Kowalko, 2020), leading to many evolutionary outputs such as the enhancing of sensory structures to navigate in the dark (Yoshizawa et al., 2010), or metabolic changes to resist long starving periods (Riddle et al., 2018; Xiong et al., 2022).

In spite of its progression and consolidation as a study model, almost nothing is known about the interactions of *Astyanax mexicanus* in nature. The studies derived from wild populations have addressed aspects of the diet (Espinasa et al., 2017), microbiome (Ornelas-García et al., 2018), circadian rhythms (Beale et al., 2013), olfactory responses (Blin et al., 2020), and parasites (Peuß et al., 2020; Santacruz et al., 2020b), covering only a few caves. The surface-dwelling *A. mexicanus* harbors a large macroparasite diversity, with ancient and highly host-specific parasite associations, mainly with helminths, the most common parasitic group, that includes trematodes, monogeneans, acantocephalans and nematodes (Pérez-Ponce de León & Choudhury, 2005; Santacruz et al., 2020a; b).

Moreover, previous studies have shown that parasites can select host adaptive traits (Hoste, 2001; Binning et al., 2017; Nadler et al., 2021), maintaining or eroding the differences in host contact hybridization zones (Theodosopoulos et al., 2019), or fueling the host divergence and speciation (Karvonen & Seehausen, 2012). Therefore, to fully understand the mechanisms operating in cavefish adaptation, it is crucial to investigate their biotic interactions, including the potential host-parasite interactions occurring in the caves.

Caves are an ideal ecological setting to test how repeated colonization of novel habitats operate in host-parasite interactions. This first requires a comprehensive understanding of parasite diversity. In this study we aim to: 1) characterize the macroparasite species diversity in 18 cave and six surface populations of *A. mexicanus*, which correspond to independent colonization lineages, 2) test the spatial rearrangement of the parasite assemblages under contrasting ecological pressures (cave and surface rivers), and 3) determine if the same parasite lineages are shared across surface and cavefish populations.

## MATERIAL AND METHODS

### Sample collection

Fish samples were sampled from 2015 to 2021 in populations along three geographical areas, Sierra de Micos (Colmena), Sierra de El Abra and Sierra de Guatemala, comprising 18 caves, and six surface populations in the proximity of the caves (Table 1). For the collection of cave specimens, permission was obtained from the competent authorities (SEMARNAT SGPA/DGVS/2438/15-16, SGPA/DGVS/05389/17, and SGPA/DGVS/1893/19). Upon capture, most of the fish were immediately euthanized in cold water with an overdose of tricain (MS-222) and preserved in absolute ethanol for later DNA extraction and parasitological screening. Other fish were scanned immediately after euthanization the same sampling day, and the voucher was preserved in ethanol. Some fish were transported to the laboratory, maintained isolated and screened for parasites several days after collection. Methods to euthanize fish were carried out in strict accordance with the American Veterinary Medical Association Guidelines for Euthanasia of Animals: 2020 edition (available at https://www.avma.org/sites/default/files/2020-02/Guidelines-on-Euthanasia-2020.pdf).

**Table 1.**
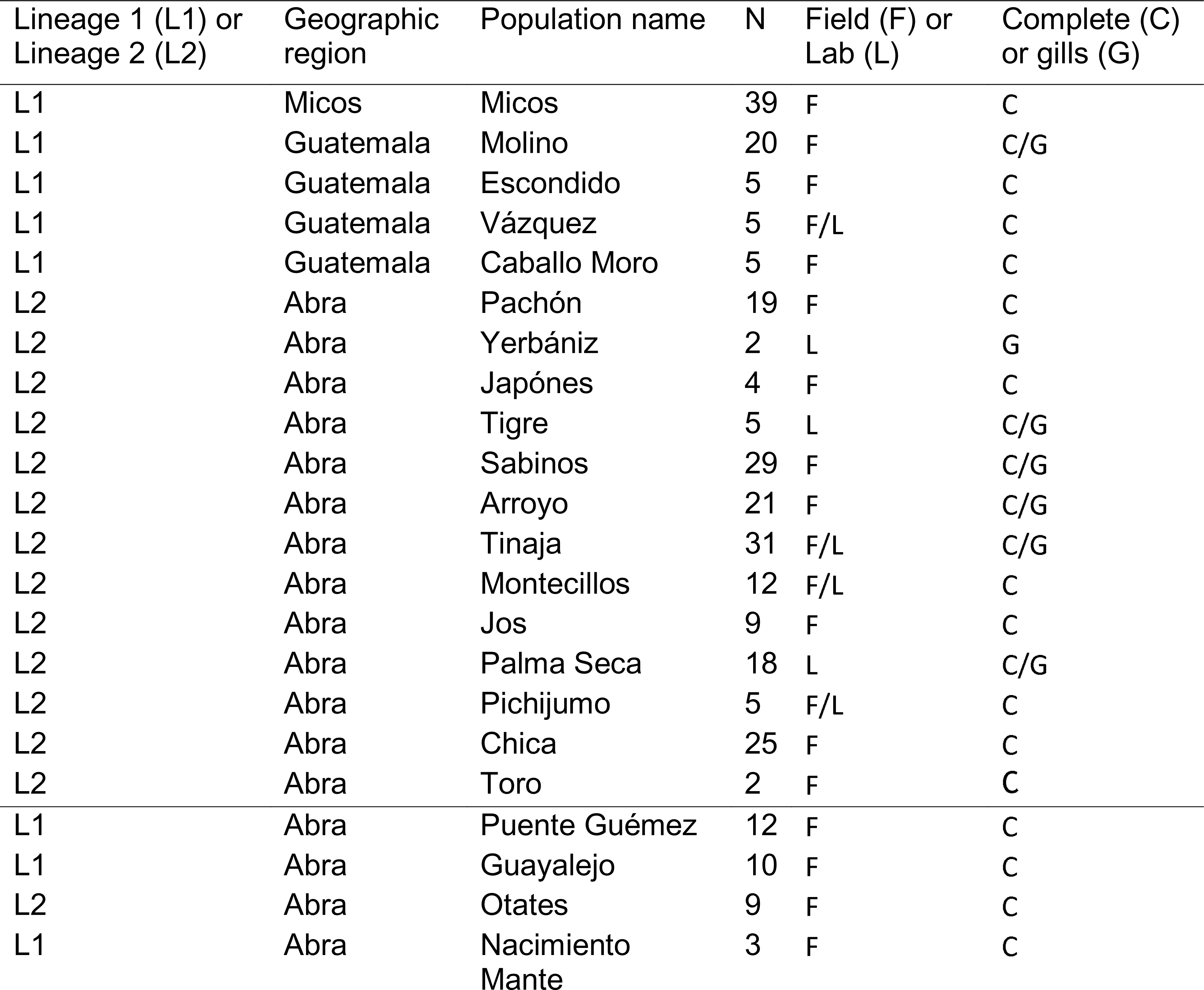
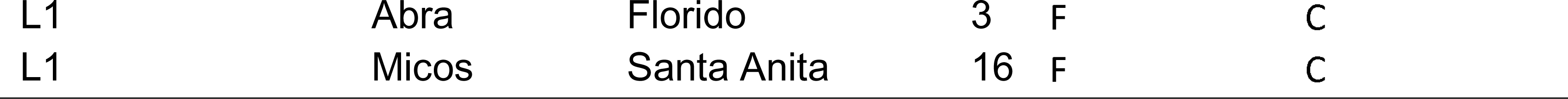
Characteristics of fish populations sampled from cave and surface populations of *Astyanax mexicanus*: lineage, geographic region, site code, number of fish individuals, fish screened in the lab or the field, and fish screened completely or only the gills.

### Parasite load

Fish analyzed on the same sampling day or stored in alcohol immediately after collection had no differences in parasitic loads. However, fish analyzed days later can suffer changes in the parasite abundance. The parasitological screening was performed under a stereomicroscope Leica Zoom 2000. Each fish was fully screened to collect macroparasites (i.e., helminths, crustaceans, mites). The scan included the skin, gills, mouth and external eyes for ectoparasites, and internal organs: internal eyes, heart, gonads, liver, gastrointestinal tract, gall bladder, spleen and abdominal cavity, for endoparasites. Parasites were removed from the host tissues with surgical needles, then washed in saline solution 6.5%, counted, and stored in molecular biology grade ethanol.

### Taxonomic identification

The identification of the parasites followed the morphological standard techniques for each group of parasites. Trematodes and monogeneans were stained with Gomori’s trichrome or Mayer’s paracarmine, whereas nematodes, acari and copepods were cleared in alcohol-glycerin (1:1) solution. Additionally, monogeneans were excised, preserving the anterior or posterior body end in ethanol. The remaining half was partially enzymatically digested to preserve sclerotized structures: haptor or male copulatory organ (MCO). Then, they were mounted in Gray-Weiss solution as permanent slides. Parasites were observed and photographed under a motorized inverted microscope Olympus IX81 equipped with differential interference contrast (DIC) optics. The ultrastructure of certain parasites was assessed by means of scanning electron microscopy (SEM) using a 10-kV Hitachi Stereoscan Model SU1510 microscope following Santacruz et al., (2020b). All parasites were identified at the highest possible taxonomic level. All the records of helminths in *A. mexicanus* in the literature were compiled from their original sources (e.g. Salgado-Maldonado et al., 2004; Salgado-Maldonado, 2006; Santacruz, 2013; Santacruz et al., 2020a; b), in addition to new records generated in this work. New species will be described in separate studies.

### Molecular analysis

For further investigation of parasite conspecificity across surface and cavefish populations, specimens fixed in 100% ethanol were individually sequenced. The molecular markers used were the mitochondrial cox1 and the nuclear genes 18S and 28S, depending on the genetic library available for each parasite group (Supporting Information, Table S1 and S2). Then, phylogenetic position was determined through Maximum likelihood using the IQ-TREE v.1.6.2 web platform (http://iqtree.cibiv.univie.ac.at/) (Nguyen et al., 2015). Genetic distances were estimated as uncorrected *p-distances* in MEGA v7 (Kumar et al., 2018).

### Data analysis

We calculated parasite species richness as the number of parasite taxa identified for each host; prevalence as the proportion of infected hosts by a given parasite species; abundance as the number of conspecific parasites in the host population; and intensity as the number of conspecific parasites in the population of infected hosts (Bush et al., 1997; Rózsa et al., 2000). Using a Wilcoxon test, we conducted paired comparisons of the parasite richness between two caves and their respective lineage’s surface population: Micos cave vs. Santa Anita river for Lineage 1, and Arroyo cave vs. Otates river for Lineage 2.

In order to evaluate the similarity of the parasite assemblages, we used the diversity values of the infracommunity. The infracommunity is defined as all the parasite species in a host at one point in time (Bush et al., 1997). With the abundance values of each parasite species, we used the Bray-Curtis dissimilarity metric as a distance measure and performed a non-metric multidimensional scaling (NMDS), using the function ‘metaMDS’ of the R package ‘vegan v1.13-1’ (Oksanen et al., 2019). In the NMDS each point represents an infracommunity. The more similar the infracommunities are, the closer the points are to each other. We used the locality, habitat (cave or surface), and geographic region as factors in the analysis. Figures were produced using the R package ‘ggplot2’ (Valero-Mora, 2016). For this analysis, we excluded individuals with incomplete data (only gills parasite data) and/or individuals that were kept in the laboratory before parasite screening. To evaluate the extent to which these differences could be explained by locality, habitat, or geographic region, we performed a PERMANOVA test.

## RESULTS

### Sampling

In total, we sampled 309 fish, 256 cavefish and 53 surface fish individuals, from 18 cave and six surface populations, respectively (Table 1, Fig. 1). We categorized the samples according to their geographical regions: Sierra de Guatemala, Sierra de El Abra and Sierra de la Colmena (Micos), and their lineage: ‘Lineage 1’ and ‘Lineage 2’.

**Figure 1.**
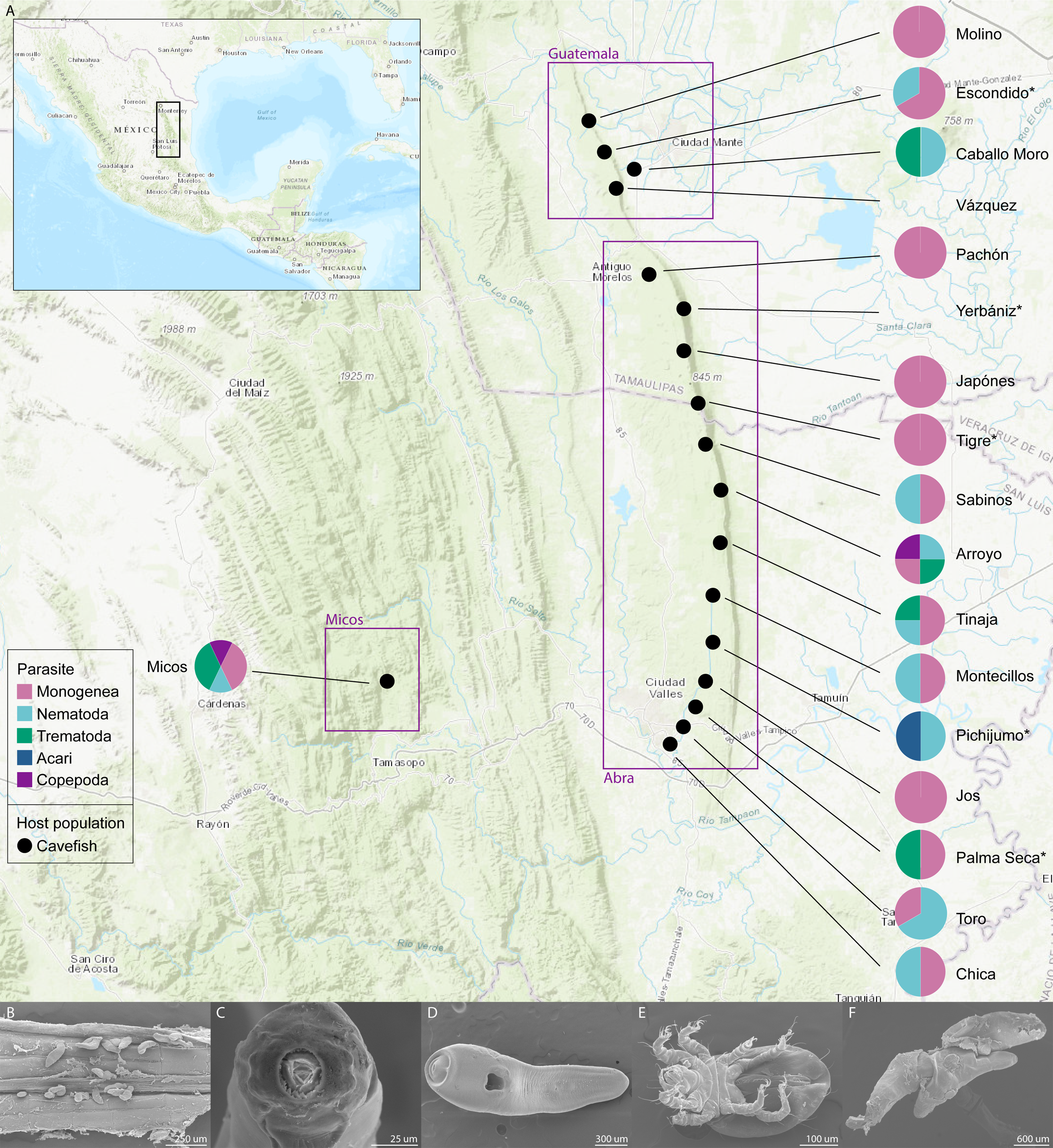
Geographical location of caves sampled in northeastern Mexico. A: The pie charts show the proportion of parasite species by taxonomic group. Data from caves Jos and Molino are only available for gill parasites. The asterisk indicates fish populations that were kept in the laboratory before parasite screening. B-F: Photomicrographs of representative parasites in the cavefish. B: *Cacatuocotyle* cf. *chajuli* monogeneans attached to the cavefish anal fin, C: anterior end of the nematode *Procamallanus neocaballeroi* Lineage 1, D: the trematode *Clinostomum* sp., E: oribatid mite, and F: the copepod *Lernaea cyprinacea*.

### Parasite diversity

We recovered 13 macroparasite taxa from cavefish populations, including endo and ecto-parasites with contrasting transmission strategies and in different life stages: larva or adults. The taxa belong to five taxonomic groups: trematodes, monogeneans, nematodes, copepods and acari (Table 2, Fig. 1). The most common parasites were monogeneans, found in 14 cavefish populations, followed by nematodes in eight, trematodes in five, acari in two and copepods in one. In Yerbaniz and Vázquez caves, no parasites were found. Except for an acari harbored by hosts from Pichijumo and Jos caves, and the unidentified trematodes from Palma Seca and Caballo Moro caves, all the parasites are a subset of the 27 taxa or lineages found across all distribution range of surface fish populations (Table 2). Five parasite taxa are shared in four or more caves: the monogeneans *Cacatuocotyle* cf. *chajuli* Mendoza-Franco, Caspeta-Mandujano & Salgado-Maldonado, 2012 and *Characithecium* cf. *costaricensis* Mendoza-Franco, Reina & Torchin, 2009, the nematodes corresponding to Lineage 1 (*sensu* Santacruz et al., 2020b) of *Procamallanus neocaballeroi* Caballero-Deloya, 1977 and *Spiroxys* sp., and the trematode *Genarchella astyanactis* Watson, 1976. One invasive species, the anchor worm, *Lernaea cyprinacea* Linnaeus, 1758 was found in a host from Micos cave.

**Table 2.**
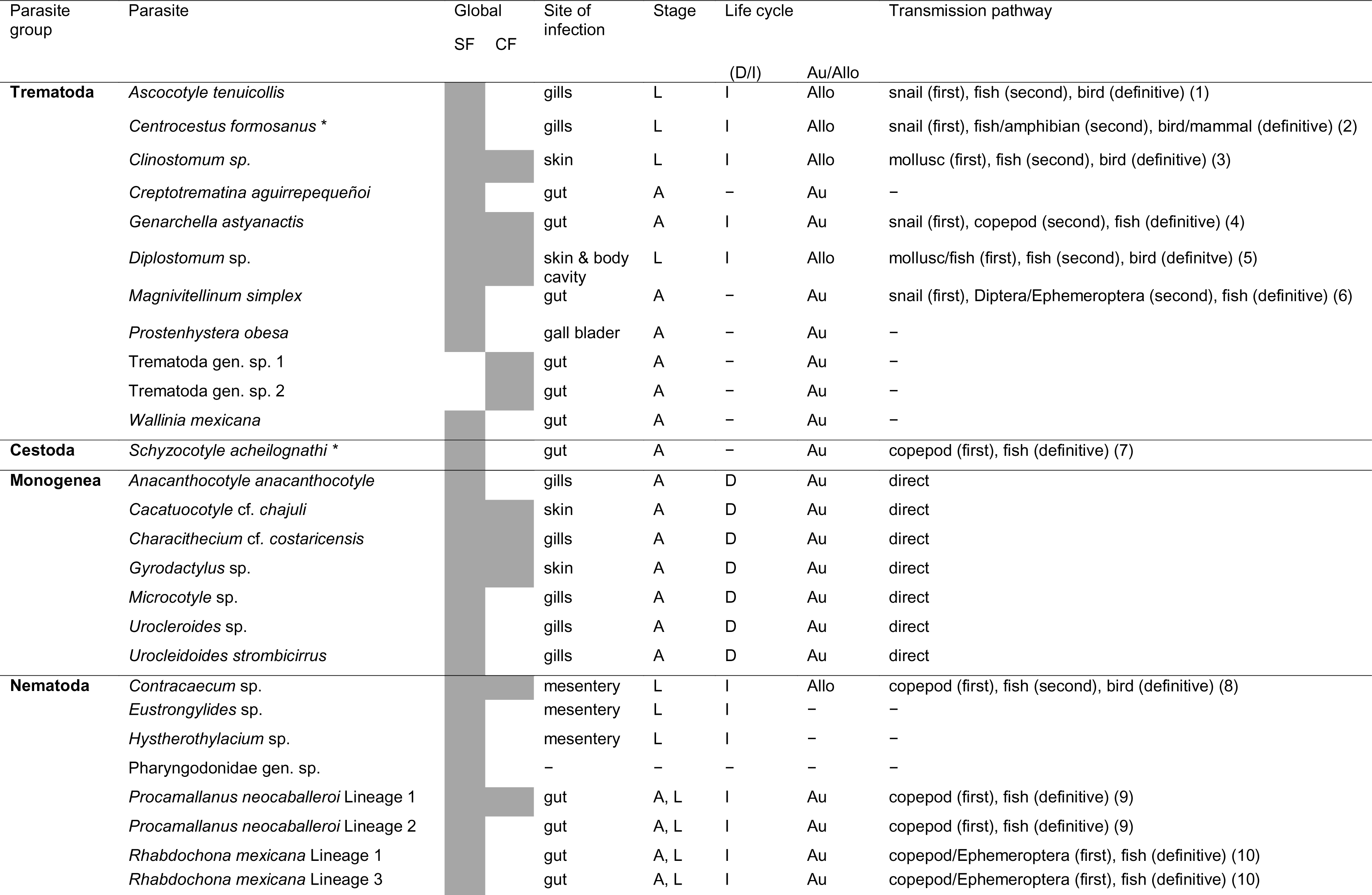
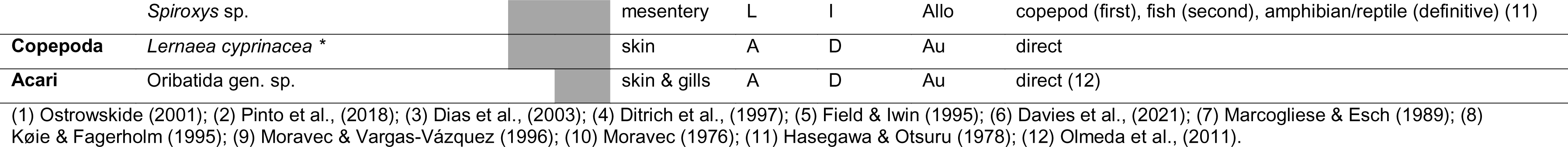
Inventory and life history traits of the parasites infecting surface and cavefish populations of *Astyanax mexicanus* across its geographical distribution. The gray cells indicate positive records in cave (CF) and/or surface (SF) fish populations. Life stage of the parasite found in the fish as larvae (L) or adult (A), type of life cycle: direct (D) or indirect (I); and autogenic (Au) or allogenic (Allo) is indicated. Asterisk indicates invasive species.

The genetic lineages of the parasites are shared across the cave and surface populations. Sequence data of some species in this study were deposited in GenBank under accession numbers: *C.* cf. *chajuli,* 28S (OQ888696–99) and COI (OQ873440–43), *C.* cf*. costaricensis*, 28S (OQ888690–95) and COI (OQ884019–22), *Spiroxys* sp., COI (OQ884015–18) and *G. astyanactis* COI (OQ873428).

### Infection patterns

Per-cavefish parasite richness varied from 0 to 4 taxa, with a mean of 0.93 taxa per host (SD= 0.95, n= 251), a maximum reached in individuals from Micos cave. Parasite species richness of surface populations ranged between 0 and 3 taxa, with a mean of 0.72 (SD= 1.18, n= 56) (Table 3, Fig. 2). The prevalence, abundance and intensity of each parasite taxa were calculated, excluding fish that were kept at the laboratory before the screening or incompletely screened for parasites (e.g., only gills). The dataset contained 183 cavefish individuals, 115 (62.8%) were infected by at least one parasite. The parasite with the highest prevalence (100%), was the gill monogenean *C*. cf. *costaricensis* in Pachón and Toro caves, followed by the nematode *Spiroxys* sp. (81%) in Micos cave (Fig. 3). The intensity and the abundance of infection were heterogenous for all the parasite taxa (Table 3). The greatest abundance and intensity for a single parasite species was displayed by *C*. cf. *chajuli* infecting fish from Toro cave, with 55 to 87 monogeneans in a single fish.

**Figure 2.**
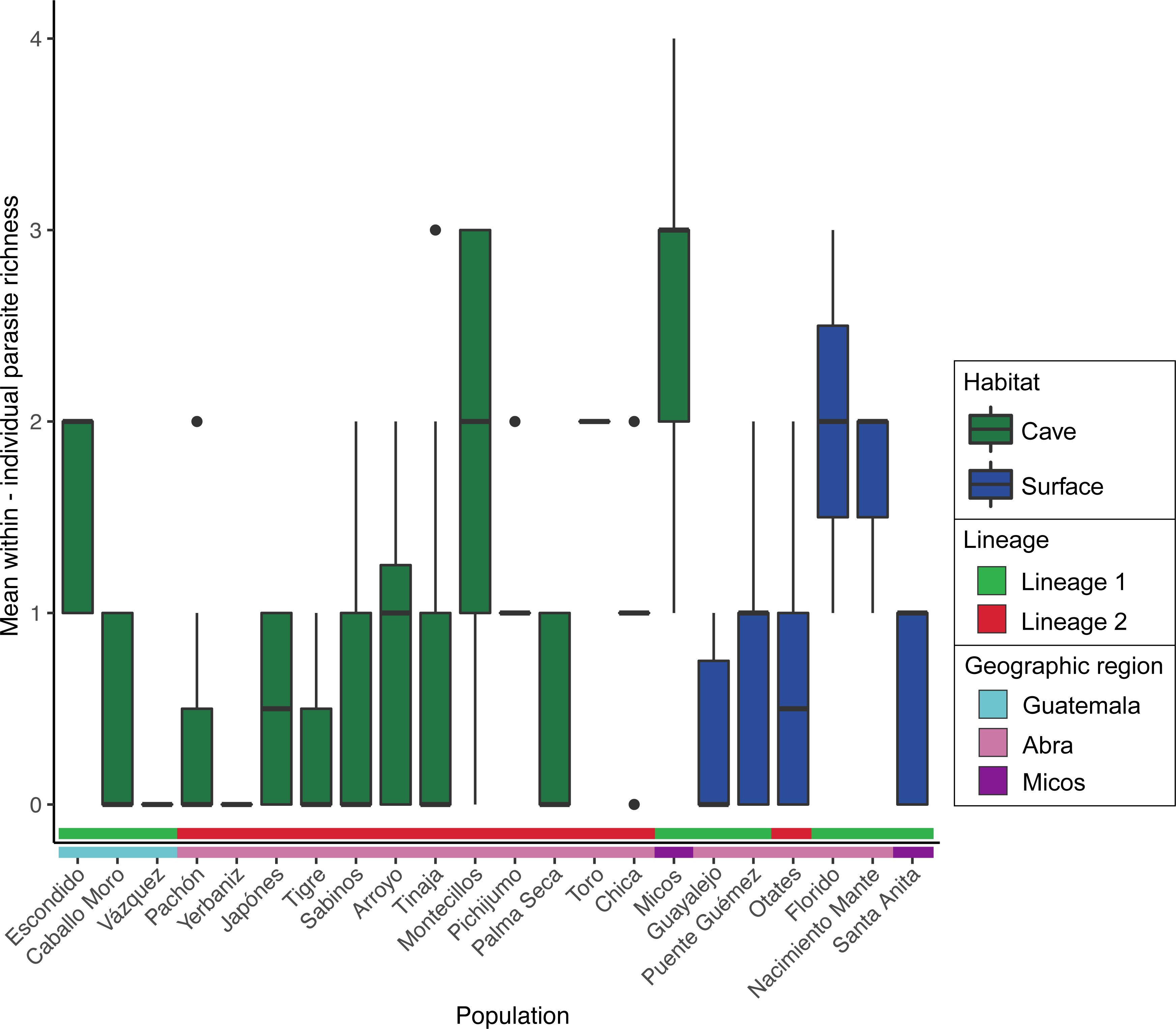
Mean individual parasite richness. Colors represent habitat, geographic region and lineage classification.

**Figure 3.**
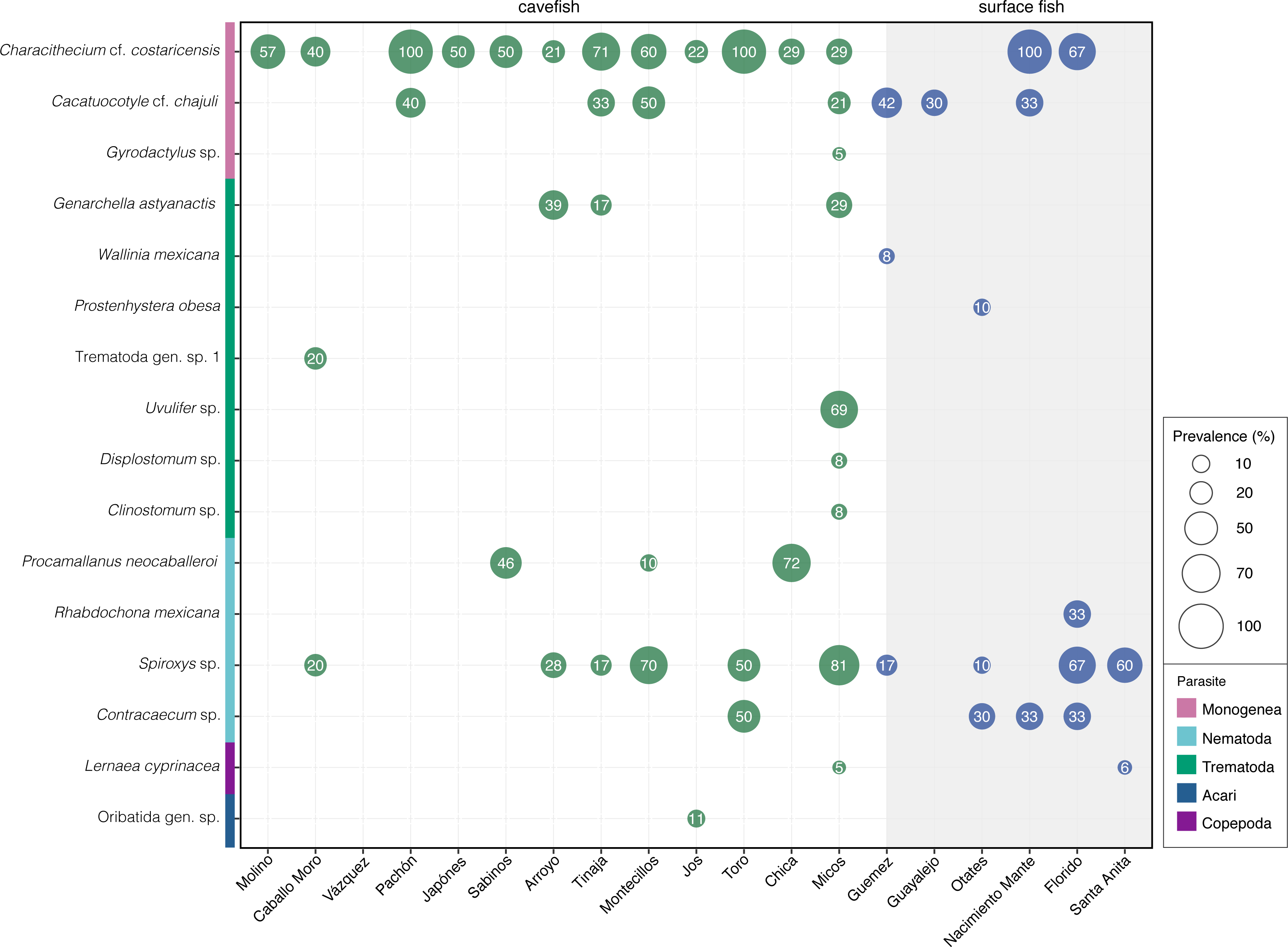
Parasite prevalence in cave and surface fish. Parasite prevalence varies among host populations and habitat. Shaded in gray the sampled surface fish populations.

**Table 3.**
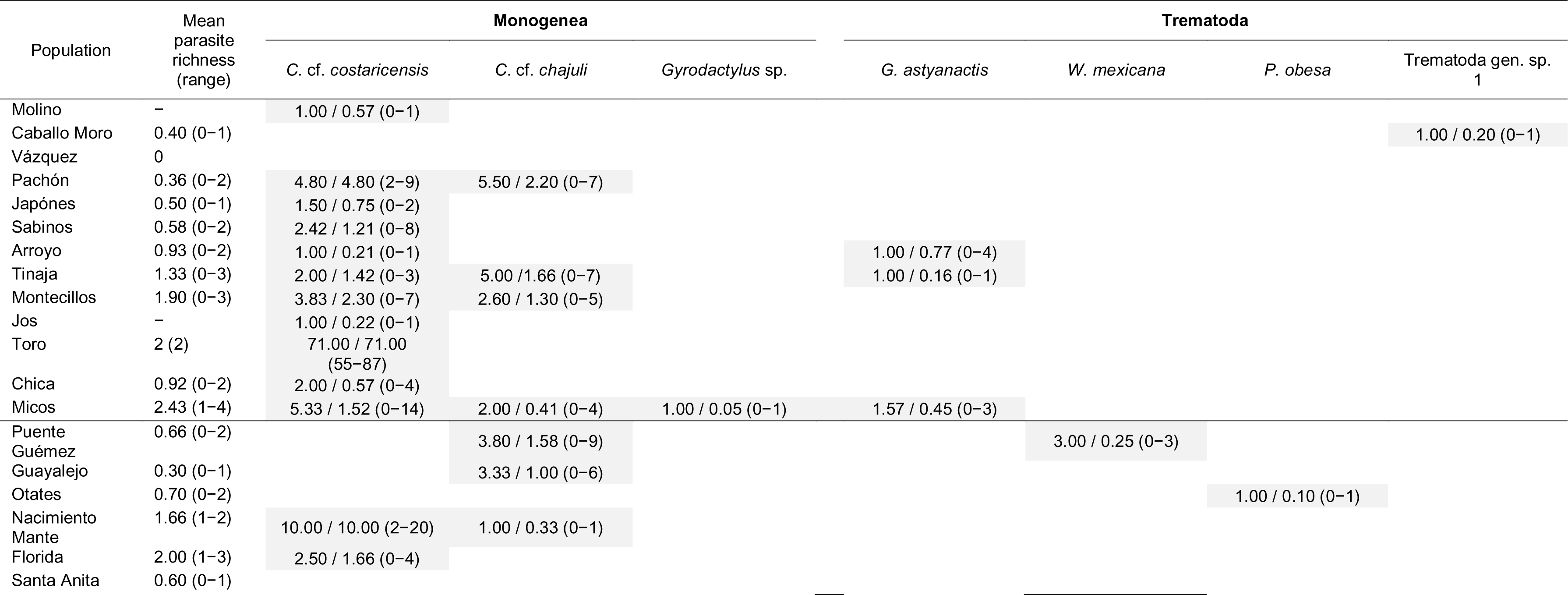
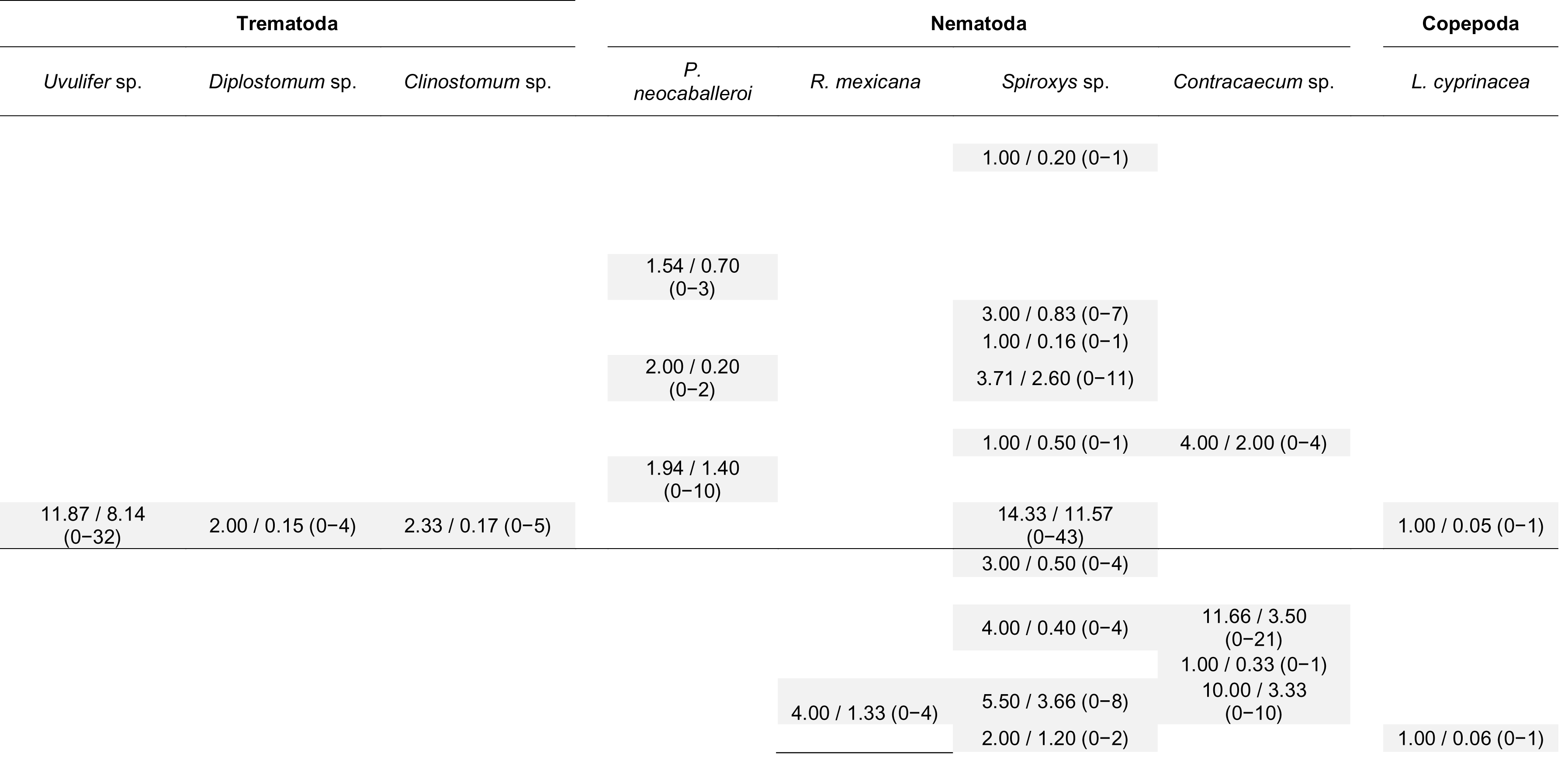
Mean parasite richness, mean intensity/mean abundance (abundance range) in cave and surface fish populations of *Astyanax mexicanus*.

The paired comparisons between cave and surface hosts showed differences in the infection profiles, the two cavefish populations considered in the analysis had higher parasite species richness than their sister surface population from the same lineage (Fig. 4). Parasite richness was significantly higher in Micos cave vs Santa Anita river (Wilcoxon test, *p<0.001*), with a total of nine parasite taxa in the cave, and a subset of two of the nine taxa were found in Santa Anita surface population. We also found significant differences between Arroyo cave vs Otates river (Wilcoxon test, *p=* 0.03); *Spiroxys* sp. was the sole species shared in both sites, and Otates river population harbored the trematode *Prostenhystera obesa* (Diesing, 1850) not found in other cave population.

**Figure 4.**
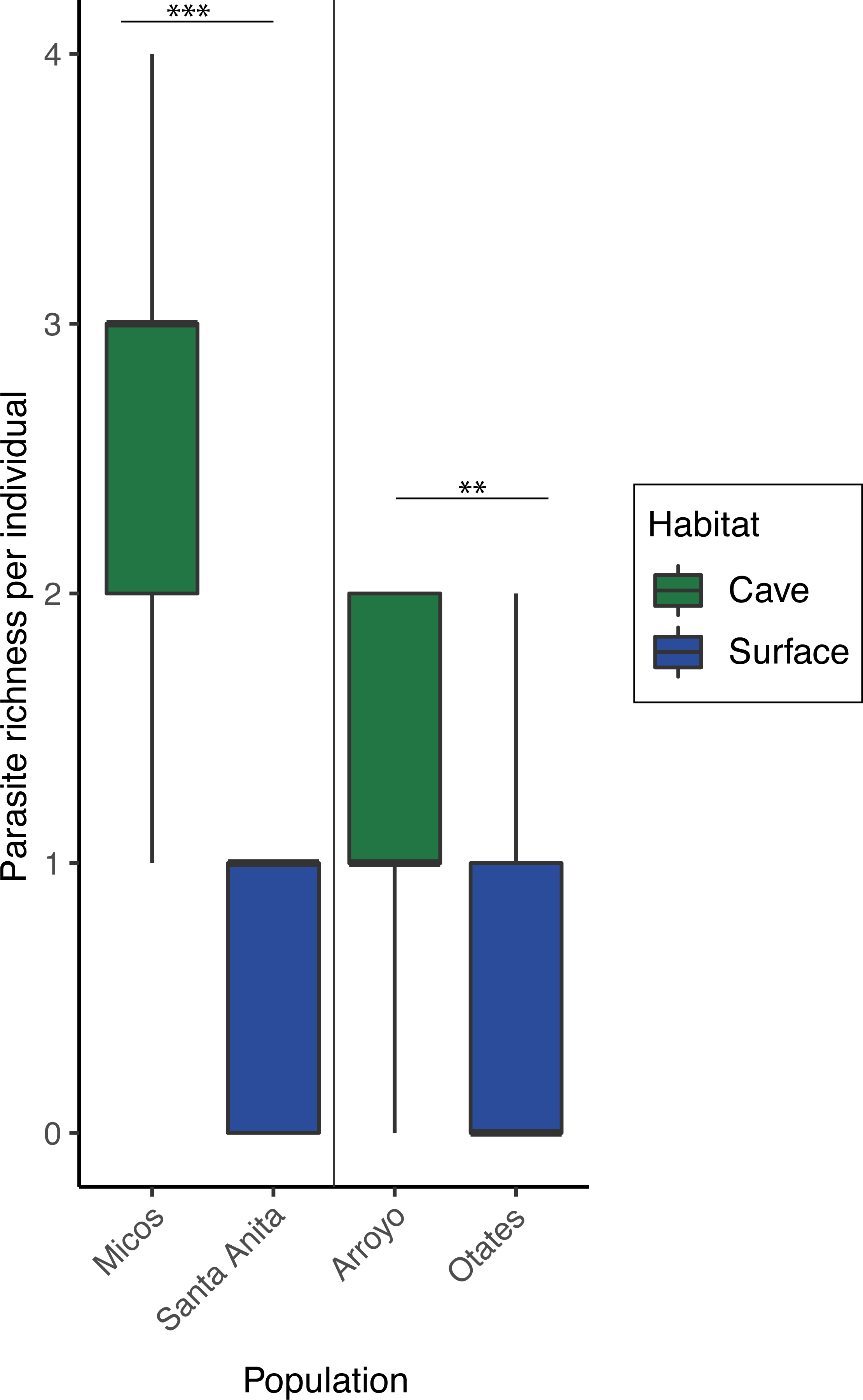
Per-fish parasite richness varies significantly between paired cave and surface populations.

### Spatial variation in parasite communities

We analyzed 125 infracommunities, 11 from caves and six from surface fish populations. The NDMS analysis showed parasite communities differentiated by population (PERMANOVA*, p=* 0.001); individuals from Micos cave were the most differentiated (Fig. 5A). The habitat (cave or surface) also had a significant effect on parasite composition (PERMANOVA*, p=* 0.002) (Fig. 5B). The strongest clusters were according to geographic region (PERMANOVA*, p=* 0.001) (Fig. 5C).

**Figure 5.**
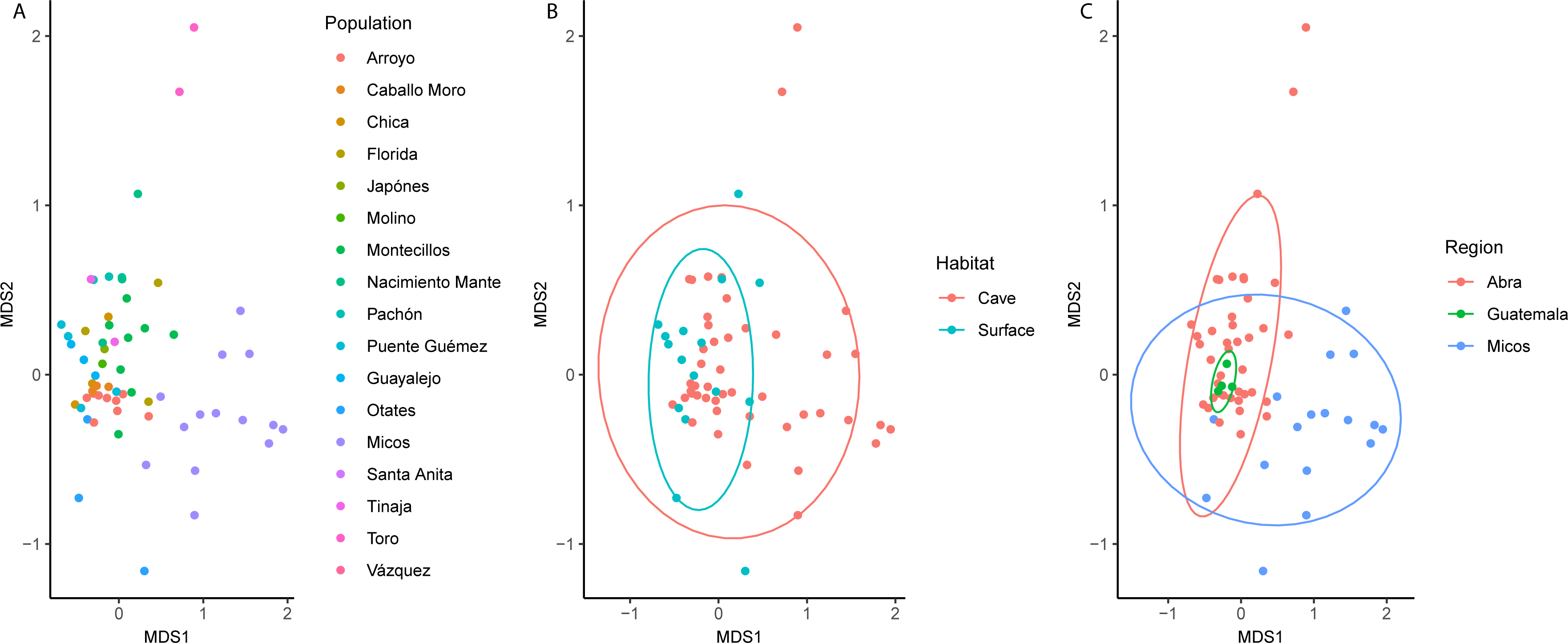
Non-metric multidimensional scaling (NMDS) plot of 125 infracommunities based on Bray-Curtis distance. Each point represents an infracommunity (all the parasites within a host). Colors indicate host population (A); habitat (B); or geographical region (C). Stress: 0.100858.

## DISCUSSION

The colonization of a new environment involves changes in the host-parasite interaction dynamics. Many parasites colonize new habitats through host-mediated dispersal, which involves a trade-off between life-history and the evolutionary pressures acting on dispersal traits (Perkins et al., 2013). Hence, during colonization the host may lose parasites generating a mosaic of parasite assemblages across landscapes (Hoberg et al., 2012).

Here, we explored the parasite diversity in cavefish populations and compared it with surface fish populations. Our study showed that parasites are very common in the caves, forming heterogeneous parasite communities across the cavefish populations, suggesting that the parasites have managed to colonize these environments together with their hosts, and persist after niche change.

In the cavefish populations examined here we identified 13 parasite taxa from distantly related taxonomic groups. Throughout the geographical distribution of surface fish, our data suggests a high diversity of parasites compared to other fish species in the same area. The cavefish harbors a subset of the global parasite diversity known for the surface *A. mexicanus*, sharing 10 of the 27 parasite taxa.

The most common cavefish parasites belong to the ‘core parasite fauna’ of Middle American characids (Pérez-Ponce de León & Choudhury, 2005). Here, based on their prevalence and abundance we propose a ‘cavefish core parasite fauna’ which includes a trematode, *G. astyanactis*, the monogeneans *C*. cf. *chajuli* and *C*. cf. *costaricensis* and the nematodes *P. neocaballeroi* and *Spiroxys* sp. We found these parasites in four or more caves, indicating that their presence could be moderately deterministic: intrinsic parasite traits may elude the same ecological filters, allowing repeated cave colonization.

Monogeneans have a direct transmission, which only requires the fish to complete its life-cycle, hence, monogeneans are the most likely candidates for co-invading the caves along with their hosts. Indeed, in our study the monogeneans have been successful colonizers, since they were found in 14 of the 18 caves analyzed. Given the low frequency of monogeneans in surface populations, it seems likely that they have experienced repeated cave invasions, increasing the likelihood of infected individuals entering the caves, and allowing the establishment of the cavefish-monogenean interaction.

In contrast, the remaining members of the ‘cavefish core parasite fauna’ display a trophic transmission strategy; some of them mature and reproduce in the fish whereas others require the fish to be eaten by a predator for the parasite to complete its life-cycle. This scenario implies that other organisms suitable as intermediate hosts also colonized the caves, giving rise to interaction networks occurring in perpetual darkness. For *G. astyanactis* and *P. neocaballeroi*, both infect copepods that are eaten by the fish (Moravec & Vargas-Vázquez, 1996; Ditrich et al., 1997). Instead, *Spiroxys* sp. found as a larva in fish, can mature in amphibians and reptiles (e.g. Li et al., 2014). That is, trophically-transmitted parasites are a good proxy for host diet and host predators predictions (Johansen et al., 2019; Leung & Koprivnikar, 2019). In cavefish from Pachón it has been documented a diet including copepods, ostracods and isopods in non-adult fish stages (Espinasa et al., 2017), which is a diet similar to our own observations in adult cavefish from different caves (Santacruz pers. obs). Regarding predators, it has been proposed they are greatly reduced in numbers in the caves, linked to the cavefish behavioral syndrome that includes vibration attraction response (VAB) and loss of schooling (Yoshizawa et al., 2010; Kowalko et al., 2013). Therefore, one possible alternative is that immature *Spiroxys* sp. somehow enter the caves representing a dead-end for the parasite.

The presence of the anchor worm *L. cyprinacea* in cavefish from Micos cave is noteworthy. The copepod is considered an invasive species (Narciso et al., 2019; Zhu et al., 2020). Furthermore, this copepod causes intense inflammation at the site of attachment leading to secondary infections (Salinas et al., 2019). The parasite has been usually co-introduced with carp across the world (Steckler & Yanong, 2012). Although we did not find carp in the cave, the parasites could have entered during flooding events with surface fish already infected. Another rare association that requires further scrutiny is that of the mites infecting the gills and skin of fish from Pichijumo and Jos caves, not yet taxonomically determined. This parasite has not been documented in surface fish, although in general oribatid mites have rarely been reported as parasites (Olmeda et al., 2011; Santacruz et al., 2022).

The heterogeneous nature of parasite communities across the caves is expected given the different scale-specific effects (e.g. host immunity, diet, sex) (see Poulin & Valtonen, 2002; Bolnick et al., 2020a; b). Numerous studies have shown that exposition to different parasites may result in local adaptations; for instance, differences in resistance or susceptibility in sticklebacks as a result of a parasite-mediated selection on the immune response and/or the mate choice (Scharsack et al., 2007; Eizaguirre et al., 2012; Milinski, 2014; Stutz et al., 2014, 2015). In our paired comparisons between cave and surface parasite infections, we uncovered differences between both environments, with a tendency for higher levels of infection and aggregation in the caves, and no sign of parallel levels of infections in the same host ecomorphotype. This difference could lead to divergence in the host immune response across the cave populations. For instance, the population of Pachón cave shows a more sensitive immune response than some surface fish (Peuß et al., 2020), confirming that local adaptations might partially explain why cavefish are more parasitized in particular populations. Additionally, cavefish populations lack some ecological pressures, including predation or interspecific competition, which may allow individuals to tolerate greater parasitic loads. Moreover, since the caves show contrasting abiotic variations compared to those of the surface rivers (Ornelas-García et al., 2018), it leads to the question of how such abiotic differences are shaping host-parasite interactions. For instance, water temperature preferences in the cavefish (see Tabin et al., 2018) have been linked to contrasting parasite infections in other fishes (also see Karvonen et al., 2013).

The parasite taxa we analyzed genetically demonstrated that the same lineages are shared between cave and surface populations. The result leads to multiple explanations. For instance, it might be possible that niche change has not -yet-promoted the parasite divergence, as the parasites recently entered the caves, resembling the recent dispersion of their host (see Herman *et al*., 2018), that might be there are subterranean connections between cave populations (as suggested by Bradic et al., 2012), or that in some caves, flooding events maintain gene flow between parasite populations. Additionally, the connectivity between the surface and the caves could also be maintaining the lack of differentiation between the surface and cave parasite lineages (see Moran et al. 2022). All the above are not mutually exclusive phenomena. A caveat in our study is that our analysis is limited to a few genetic markers, and few parasite individuals. It would be valuable to analyze other genomic regions in order to have a detailed picture of the parasite evolutionary history and to test if they recover the same patterns and time frame of colonization as their hosts. The prime candidates for further investigation regarding host-parasite phylogenetic congruence and coevolution are the monogeneans, given their close and predominant association with the cavefish, as well as elevated values of prevalence, abundance, and intensity of infection, and the fact that their transmission route is direct. Both species of monogeneans found in the cavefish, i.e., *C*. cf. *chajuli* and *C*. cf. *costaricensis* are considered species complexes, which will be described elsewhere (Santacruz pers. comm.).

Previous studies about parasites in troglobitic organisms are very rare. For instance, in the blind catfish (Moravec & Huffman, 1988), or in cave animals such as guppies and salamanders (e.g. Dyer & Peck, 1975; Tobler et al., 2014). *Astyanax mexicanus* is a study model in eco-evo-devo (Casane & Rétaux, 2016; Krishnan & Rohner, 2017; Jeffery, 2020), hence the knowledge of its ecological interactions in its natural environment are fundamental for understanding the mechanisms driving its adaptation to the caves. The variation of parasite infections across the caves offers an invaluable opportunity to test the role of parasites in the contrasting physiological, morphological, metabolic and behavioral adaptive changes that could have allowed *A. mexicanus* to colonize an extreme environment.

## CONCLUSIONS

We studied the parasites infecting the cavefish, providing the first parasitological records for more than half of their known populations. Our results indicate: (1) great parasite diversity composed by distantly related parasites, with contrasting life histories, (2) most of the parasites found in the caves are a subset of the parasite diversity known for surface-dwelling populations, (3) some parasite species are more frequent in the caves than in surface (i.e. monogeneans), making up the ‘cavefish core parasite fauna’, (4) the parasite communities vary across cavefish populations and, (5) cave and surface populations differ in their infections. More caves remain to be explored to test if the patterns found in our study are still recovered. Nonetheless, our parasite inventory opens numerous avenues and questions regarding how a host-parasite interaction is shaped in a cave extreme environment. Further investigation including the abiotic variables in the caves, together with the parasite evolutionary history could give us insights into the factors driving the parasite diversification process in the caves.

## DATA AVAILABILITY

The fish were deposited in Colección Nacional de Peces, Instituto de Biología, UNAM. The parasitological material was deposited in the national collections of the National Autonomous University of Mexico (UNAM), Mexico City: Colección Nacional de Helmintos (CNHE) and Colección Nacional de Ácaros (CNAC).

## SUPPLEMENTARY DATA

Supplementary data to this article can be found online.

## AUTHOR’S CONTRIBUTIONS

A.S. conceived the study, contributed to fish sampling, performed the fish parasitological screening, identified the parasites, performed the experiments, analyzed the data, prepared figures and/or tables, wrote the paper.

D.H.M. contributed to fish sampling and fish parasitological screening, reviewed and edited the manuscript.

R.M.G. contributed to fish sampling.

G.P.P.L. contributed to parasite identification, contributed reagents/materials, reviewed and edited the manuscript.

P.O.G. conceived the study, performed the fish sampling, contributed reagents/materials, reviewed and edited the manuscript.

All authors read and approved the final version of the manuscript.

## ACKNOWLEDGMENTS

We thank Edgar Mendoza Franco and Carlos Mendoza Palmero for their help with monogenenan identification. We also want to thank Carlos Lozano for his kind suggestions on an earlier version of this manuscript. We also thank to Miriam Miranda Gamboa, Carlos Pedraza Lara, Ulises Ribeiro, and Angeles Verde for their help in sample collection. Berenit Mendoza from the LaNaBio for her support with the SEM photomicrographs. Andrea Jiménez, Laura Márquez and Nelly López also from the LaNaBio for their help in the molecular work. We thank the anonymous reviewers for their invaluable comments.

## SUPPORTING INFORMATION

**Table S1.** Primers used for PCR and sequencing of parasites in this study.

**Table S2.** Primer combination and PCR conditions.

## References

Beale A, Guibal C, Tamai TK, Klotz L, Cowen S, Peyric E, et al. 2013. Circadian rhythms in Mexican blind cavefish *Astyanax mexicanus* in the lab and in the field. Nat. Commun. 4.

Best A, Ashby B, White A, Bowers R, Buckling A, Koskella B, et al. 2017. Host-parasite fluctuating selection in the absence of specificity. Proc. R. Soc. B Biol. Sci. 284: 20171615.

Binning SA, Shaw AK, Roche DG. 2017. Parasites and host performance: Incorporating infection into our understanding of animal movement. Integr. Comp. Biol. 57: 267– 280.

Blin M, Fumey J, Lejeune C, Policarpo M, Leclercq J, Père S, et al. 2020. Diversity of olfactory responses and skills in *Astyanax mexicanus* cavefish populations inhabiting different caves. Diversity 12: 1–21.

Bolnick DI, Resetarits EJ, Ballare K, Stuart YE, Stutz WE. 2020a. Host patch traits have scale-dependent effects on diversity in a stickleback parasite metacommunity. Ecography 43: 990–1002.

Bolnick DI, Resetarits EJ, Ballare K, Stuart YE, Stutz WE. 2020b. Scale-dependent effects of host patch traits on species composition in a stickleback parasite metacommunity. Ecology 101: e03181.

Bradic M, Beerli P, García-De Leán FJ, Esquivel-Bobadilla S, Borowsky RL. 2012. Gene flow and population structure in the Mexican blind cavefish complex (*Astyanax mexicanus*). BMC Evol. Biol. 12: 1–17.

Brunner FS, Anaya-Rojas JM, Matthews B, Eizaguirre C. 2017. Experimental evidence that parasites drive eco-evolutionary feedbacks. Proc. Natl. Acad. Sci. U. S. A. 114: 3678–3683.

Brunner FS, Eizaguirre C. 2016. Can environmental change affect host/parasite-mediated speciation? Zoology 119: 384–394.

Bush AO, Lafferty KD, Lotz JM, Shostak AW. 1997. Parasitology meets ecology on its own terms: Margolis et al. revisited. J. Parasitol. 83: 575–583.

Bush SE, Villa SM, Altuna JC, Johnson KP, Shapiro MD, Clayton DH. 2019. Host defense triggers rapid adaptive radiation in experimentally evolving parasites. Evol. Lett. 3: 120–128.

Casane D, Rétaux S. 2016. Evolutionary Genetics of the Cavefish *Astyanax mexicanus*. Adv. Genet. 95: 117–159.

Colautti RI, Ricciardi A, Grigorovich IA, MacIsaac HJ. 2004. Is invasion success explained by the enemy release hypothesis? Ecol. Lett. 7: 721–733.

Criscione CD, Blouin MS. 2004. Life cycles shape parasite evolution: comparative population genetics of salmon trematodes. Evolution 58: 198–202.

Davies D, Liquin F, Lauthier JJ, Párraga R, Saravia J, Davies C, et al. 2021. The life cycle of *Magnivitellum saltaensis* n. sp. (Digenea: Alloglossidiidae) in Salta Province, Argentina. Parasitol. Res. 120: 1233–1245.

Demandt N, Saus B, Kurvers RHJM, Krause J, Kurtz J, Scharsack JP. 2018. Parasite-infected sticklebacks increase the risk-taking behavior of uninfected group members. Proc. R. Soc. B Biol. Sci. 285: 20180956.

Dias MLGG, Eiras JC, Machado MH, Souza GTR, Pavanelli GC. 2003. The life cycle of *Clinostomum complanatum* Rudolphi, 1814 (Digenea, Clinostomidae) on the floodplain of the high Paraná river, Brazil. Parasitol Res. 89: 506–8.

Ditrich O, Scholz T, Aguirre-Macedo ML, Vargas-Vázquez J. 1997. Larval stages of trematodes from freshwater molluscs of the Yucatan Peninsula, Mexico. Folia Parasitol. 44: 109–127.

Dyer WG, Peck SB. 1975. Gastrointestinal parasites of the cave salamander, *Eurycea lucifuga* Rafinesque, from the southeastern United States. Can. J. Zool. 53: 52–54.

Eizaguirre C, Lenz TL. 2010. Major histocompatibility complex polymorphism: Dynamics and consequences of parasite-mediated local adaptation in fishes. J. Fish Biol. 77: 2023–2047.

Eizaguirre C, Lenz TL, Kalbe M, Milinski M. 2012. Rapid and adaptive evolution of MHC genes under parasite selection in experimental vertebrate populations. Nat. Commun. 3: 1–6.

Eizaguirre C, Lenz TL, Sommerfeld RD, Harrod C, Kalbe M, Milinski M. 2011. Parasite diversity, patterns of MHC II variation and olfactory based mate choice in diverging three-spined stickleback ecotypes. Evol. Ecol. 25: 605–622.

Elliot WR, Russell WH, Mitchell RW. 2019. The *Astyanax* caves of Mexico: cavefishes of Tamaulipas, San Luis Potosí and Guerrero. J. Fish Biol. 94: 205–205.

Erin NI, Benesh DP, Henrich T, Samonte IE, Jakobsen PJ, Kalbe M. 2019. Examining the role of parasites in limiting unidirectional gene flow between lake and river sticklebacks. J. Anim. Ecol. 88: 1986–1997.

Espinasa L, Bonaroti N, Wong J, Pottin K, Queinnec E, Rétaux S. 2017. Contrasting feeding habits of post-larval and adult *Astyanax* cavefish. Subterr. Biol. 21: 1–17.

Espinasa L, Legendre L, Fumey J, Blin M, Rétaux S, Espinasa M. 2018. A new cave locality for *Astyanax* cavefish in Sierra de El Abra, Mexico. Subterr. Biol. 26: 39–53.

Espinasa L, Ornelas-García CP, Legendre L, Rétaux S, Best A, Gamboa-Miranda R, et al. 2020. Discovery of Two New *Astyanax* Cavefish Localities Leads to Further Understanding of the Species Biogeography. Diversity 12: 368.

Field JS, Irwin WB. 1995. Life-cycle description and comparison of *Displostomum spathaceum* (Rudolphi, 1819) and *D. pseudobaeri* (Razmaskin & Andrejak, 1978) from rainbow trout (*Oncorhynchus mykiss* Walbaum) maintained in identical hosts. Parasitol. Res. 81: 505–517.

Garduño-Sánchez M, Hernandez-Lozano J, Moran R, Miranda-Gamboa R, Gross J, Rohner N, et al. 2022. Phylogeographic relationships and morphological evolution between cave and surface *Astyanax mexicanus* populations (De Fillipi 1853) (Actinopterygii, Characidae). (In review).

Gross JB, Powers AK. 2020. A Natural Animal Model System of Craniofacial Anomalies: The Blind Mexican Cavefish. Anat. Rec. 303: 24–29.

Hasegawa H, Otsuru M. 1978. Notes on the Life Cycle of *Spiroxys japonica* Morishita, 1926 (Nematoda: Gnathostomatidae). Jap. J. Parasit. 2: 113–122.

Herman A, Brandvain Y, Weagley J, Jeffery WR, Keene AC, Kono TJY, et al. 2018. The role of gene flow in rapid and repeated evolution of cave-related traits in Mexican tetra, *Astyanax mexicanus*. Mol. Ecol. 27: 4397–4416.

Hoberg EP, Galbreath KE, Cook JA, Kutz SJ, Polley L. 2012. Northern host-parasite assemblages. History and biogeography on the borderlands of episodic climate and environmental transition. Adv. Parasitol. 79: 1–97.

Hoste H. 2001. Adaptive physiological processes in the host during gastrointestinal parasitism. Int. J. Parasitol. 31: 231–244.

Hyacinthe C, Attia J, Rétaux S. 2019. Evolution of acoustic communication in blind cavefish. Nat. Commun. 10: 1–12.

Jeffery WR. 2020. *Astyanax* surface and cave fish morphs. Evodevo 11: 14.

Johansen IB, Henriksen EH, Shaw JC, Mayer I, Amundsen PA, Øverli. 2019. Contrasting associations between breeding coloration and parasitism of male Arctic charr relate to parasite species and life cycle stage. Sci. Rep. 9: 1–9.

Jolles JW, Mazué GPF, Davidson J, Behrmann-Godel J, Couzin ID. 2020. *Schistocephalus* parasite infection alters sticklebacks’ movement ability and thereby shapes social interactions. Sci. Rep. 10: 12282.

Karvonen A, Kristjánsson BK, Skúlason S, Lanki M, Rellstab C, Jokela J. 2013. Water temperature, not fish morph, determines parasite infections of sympatric Icelandic threespine sticklebacks (*Gasterosteus aculeatus*). Ecol. Evol. 3: 1507.

Karvonen A, Seehausen O. 2012. The role of parasitism in adaptive radiations-when might parasites promote and when might they constrain ecological speciation? *Int*. J. Ecol. 2012.

Koie M, Fagerholm HP. 1995. The life cycle of *Contracaecum osculatum* (Rudolphi, 1802) sensu stricto (Nematoda, Ascaridoidea, Anisakidae) in view of experimental infections. Parasitol. Res. 81: 481–9.

Kowalko J. 2020. Utilizing the blind cavefish *Astyanax mexicanus* to understand the genetic basis of behavioral evolution. J. Exp. Biol. 223.

Kowalko JE, Rohner N, Rompani SB, Peterson BK, Linden TA, Yoshizawa M, et al. 2013.Loss of schooling behavior in cavefish through sight-dependent and sight-independent mechanisms. Curr. Biol. 23: 1874–1883.

Krishnan J, Rohner N. 2017. Cavefish and the basis for eye loss. Philos. Trans. R. Soc. Lond. B. Biol. Sci. 372.

Kumar S, Stecher G, Li M, Knyaz C, Tamura K. 2018. MEGA X: Molecular Evolutionary Genetics Analysis across computing platforms. Mol. Biol. Evol. 35: 1547–1549.

Leung TLF, Koprivnikar J. 2019. Your infections are what you eat: How host ecology shapes the helminth parasite communities of lizards. J. Anim. Ecol. 88: 416–426.

Li L, Hasegawa H, Roca V, Xu Z, Guo YN, Sato A, et al. 2014. Morphology, ultrastructure and molecular characterisation of *Spiroxys japonica* Morishita, 1926 (Spirurida: Gnathostomatidae) from *Pelophylax nigromaculatus* (Hallowell) (Amphibia: Ranidae). Parasitol. Res. 113: 893–901.

Lymbery AJ. 2015. Niche construction: Evolutionary implications for parasites and hosts. Trends Parasitol. 31: 134–141.

Marcogliese DJ, Esch GW. 1989. Experimental and Natural Infection of Planktonic and Benthic Copepods by the Asian Tapeworm, *Bothriocephalus acheilognathi*. Proc. Helminthol. Soc. Wash. 56: 151–155.

Mikheev VN, Pasternak AF, Taskinen J, Valtonen TE. 2013. Grouping facilitates avoidance of parasites by fish. Parasites and Vectors 6: 301.

Milinski M. 2014. Arms races, ornaments and fragrant genes: The dilemma of mate choice in fishes. Neurosci. Biobehav. Rev. 46: 567–572.

Miranda-Gamboa R, Espinasa L, Verde-Ramírez MA, Hernández-Lozano J, Lacaille JL, Espinasa M, et al. 2023. A new cave population of *Astyanax mexicanus* from Northern Sierra de El Abra, Tamaulipas, Mexico. Subterr. Biol. 45: 95–117.

Moran RL, Jaggard JB, Roback EY, Kenzior A, Rohner N, Kowalko JE, et al. 2022. Hybridization underlies localized trait evolution in cavefish. iScience 25: 103778.

Moran RL, Richards EJ, Ornelas-García CP, Gross JB, Donny A, Wiese J, et al. 2023. Selection-driven trait loss in independently evolved cavefish populations. Nat. Commun. 14: 2557.

Moravec F. 1976. Observations on the development of *Rhabdochona phoxini* Moravec, 1968 (Nematoda: Rhabdochonidae). Folia Parasitol. 23: 309–20.

Moravec F, Huffman DG. 1988. *Rhabdochona longleyi* sp. n. (Nematoda: Rhabdochonidae) from blind catfishes, *Trogloglanis pattersoni* and *Satan eurystomus* (Ictaluridae) from the subterranean waters of Texas. Folia Parasitol. 35: 235–243.

Moravec F, Vargas-Vázquez J. 1996. The development of *Procamallanus (Spirocamallanus) neocaballeroi* (Nematoda: Camallanidae), a parasite of *Astyanax fasciatus* (Pisces) in Mexico. Folia Parasitol. 43: 61–70.

Nadler LE, Bengston E, Eliason EJ, Hassibi C, Helland-Riise SH, Johansen IB, et al. 2021. A brain-infecting parasite impacts host metabolism both during exposure and after infection is established. Funct. Ecol. 35: 105–116.

Narciso RB, Acosta AA, Nobile AB, de Lima FP, Freitas-Souza D, da Silva RJ. 2019. *Lernaea cyprinacea* (Copepoda: Lernaeidae) in *Piabarchus stramineus* (Characiformes: Characidae) from the Taquari River, São Paulo State, Brazil. Biologia. 74: 1171–1179.

Nguyen LT, Schmidt HA, Von Haeseler A, Minh BQ. 2015. IQ-TREE: A fast and effective stochastic algorithm for estimating maximum-likelihood phylogenies. Mol. Biol. Evol. 32: 268–274.

Oksanen J, Blanchet FG, Friendly M, Kindt R, Legendre P, McGlinn D, et al. 2019. vegan: Community Ecology Package.

Olmeda AS, Blanco MM, Pérez-Sánchez JL, Luzón M, Villarroel M, Gibello A. 2011. Occurrence of the oribatid mite *Trhypochthoniellus longisetus longisetus* (Acari: Trhypochthoniidae) on tilapia *Oreochromis niloticus*. Dis. Aquat. Organ. 94: 77–81.

Ornelas-García CP, Domínguez-Domínguez O, Doadrio I. 2008. Evolutionary history of the fish genus *Astyanax* Baird & Girard (1854) (Actinopterygii, Characidae) in Mesoamerica reveals multiple morphological homoplasies. BMC Evol. Biol. 8: 1–17.

Ornelas-García P, Pajares S, Sosa-Jiménez VM, Rétaux S, Miranda-Gamboa RA. 2018. Microbiome differences between river-dwelling and cave-adapted populations of the fish *Astyanax mexicanus* (De Filippi, 1853). PeerJ 6, e5906.

Ostrowskide NM. 2001. Life cycles of two new sibling species of *Ascocotyle* (*Ascocotyle*) (Digenea, Heterophyidae) in the Neotropical Region. Acta Parasitol. 46: 119–129.

Pérez-Ponce de León G, Choudhury A. 2005. Biogeography of helminth parasites of freshwater fishes in Mexico: The search for patterns and processes. J. Biogeogr. 32: 645–659.

Perkins AT, Phillips BL, Baskett ML, Hastings A. 2013. Evolution of dispersal and life history interact to drive accelerating spread of an invasive species. Ecol. Lett. 16: 1079–1087.

Peuß R, Box AC, Chen S, Wang Y, Tsuchiya D, Persons JL, et al. 2020. Adaptation to low parasite abundance affects immune investment strategy and immunopathological responses of cavefish. *Nat*. Ecol. Evol. 4: 1416–1430.

Pinto HA, Gonçalves NQ, López-Hernández D, Pulido-Murillo EA, Melo AL. 2018. The life cycle of a zoonotic parasite reassessed: Experimental infection of *Melanoides tuberculata* (Mollusca: Thiaridae) with *Centrocestus formosanus* (Trematoda: Heterophyidae). PLoS One 13, e0194161.

Poulin R, Valtonen ET. 2002. The predictability of helminth community structure in space: A comparison of fish populations from adjacent lakes. Int. J. Parasitol. 32: 1235– 1243.

Riddle MR, Aspiras AC, Gaudenz K, Peuß R, Sung JY, Martineau B, et al. 2018. Insulin resistance in cavefish as an adaptation to a nutrient-limited environment. Nature 555: 647–651.

Roose AK, Volckaert FAMM, Raeymaekers JAMM, Hablützel PI, Vanhove MPM, Deschepper P, et al. 2017. Parasite escape through trophic specialization in a species flock. J. Evol. Biol. 30: 1437–1445.

Rózsa L, Reiczigel J, Majoros G. 2000. Quantifying parasites in samples of hosts. J. Parasitol. 86: 228–232.

Salgado-Maldonado G. 2006. Checklist of helminth parasites of freshwater fishes from Mexico. Zootaxa 1324: 1–357.

Salgado-Maldonado G, Cabañas-Carranza G, Soto-Galera E, Pineda-López RF, Caspeta-Mandujano JM, Aguilar-Castellanos E, et al. 2004. Helminth parasites of freshwater fishes of the Pánuco River basin, east central Mexico. Comp. Parasitol. 71: 190–202.

Salinas ZA, Babini MS, Grenat PR, Biolé FG, Martino AL, Salas NE. 2019. Effect of parasitism of *Lernaea cyprinacea* on tadpoles of the invasive species *Lithobates catesbeianus*. Heliyon 5: e01834.

Santacruz, A. 2013. Análisis de las comunidades de peces y parásitos en la cuenca del Pánuco. Universidad Autónoma de Querétaro.

Santacruz A, Barluenga M, Pérez-Ponce de León GP. 2022. Macroparasite diversity in cichlid fish from Nicaraguan lakes, a model system for understanding host-parasite diversification and speciation. Sci. Rep. 12: 1–14.

Santacruz A, Ornelas-García CP, Pérez-Ponce de León G. 2020a. Diversity of *Rhabdochona mexicana* (Nematoda: Rhabdochonidae), a parasite of *Astyanax* spp. (Characidae) in Mexico and Guatemala, using mitochondrial and nuclear genes, with the description of a new species. J. Helminthol. 94.

Santacruz A, Ornelas-García CP, Pérez-Ponce de León G. 2020b. Incipient genetic divergence or cryptic speciation? *Procamallanus* (Nematoda) in freshwater fishes (*Astyanax*). Zool. Scr. 49: 768–778.

Scharsack JP, Kalbe M, Harrod C, Rauch G. 2007. Habitat-specific adaptation of immune responses of stickleback (*Gasterosteus aculeatus*) lake and river ecotypes. Proc. R. Soc. B Biol. Sci. 274: 1523.

Steckler N, Yanong RPE. 2012. *Lernaea* (anchorworm) *Infestations in Fish*.

Strecker U, Faúndez VH, Wilkens H. 2004. Phylogeography of surface and cave *Astyanax* (Teleostei) from Central and North America based on cytochrome b sequence data. Mol. Phylogenet. Evol. 33: 469–481.

Stutz WE, Lau OL, Bolnick DI. 2014. Contrasting patterns of phenotype-dependent parasitism within and among populations of threespine stickleback. Am. Nat. 183: 810–825.

Stutz WE, Schmerer M, Coates JL, Bolnick DI. 2015. Among-lake reciprocal transplants induce convergent expression of immune genes in threespine stickleback. Mol. Ecol. 24: 4629–4646.

Tabin JA, Aspiras A, Martineau B, Riddle M, Kowalko J, Borowsky R, et al. 2018. Temperature preference of cave and surface populations of *Astyanax mexicanus*. Dev. Biol. 441: 338–344.

Theodosopoulos AN, Hund AK, Taylor SA. 2019. Parasites and Host Species Barriers in Animal Hybrid Zones. Trends Ecol. Evol. 34: 19–30.

Tobler M, Plath M, Riesch R, Schlupp I, Grasse A, Munimanda GK, et al. 2014. Selection from parasites favours immunogenetic diversity but not divergence among locally adapted host populations. J. Evol. Biol. 27: 960–974.

Valero-Mora PM. 2016. ggplot2: Elegant Graphics for Data Analysis. J. Stat. Softw. 35.

Wegner KM, Kalbe M, Kurtz J, Reusch TBH, Milinski M. 2003. Parasite selection for immunogenetic optimality. Science 301: 1343.

Wilkens H, Strecker U. 2017. *Evolution in the Dark: Introduction*. Springer-Verlag New York, Germany.

Wolinska J, Lively CM, Spaak P. 2008. Parasites in hybridizing communities: the Red Queen again? Trends Parasitol. 24: 121–126.

Xiong S, Wang W, Kenzior A, Olsen L, Krishnan J, Persons J, et al. 2022. Enhanced lipogenesis through Pparγ helps cavefish adapt to food scarcity. Current Biology, 32: 2272–2280.

Yoshizawa M, Gorički Š, Soares D, Jeffery WR. 2010. Evolution of a behavioral shift mediated by superficial neuromasts helps cavefish find food in darkness. Curr. Biol. 20: 1631.

Zhu X, Barton DP, Wassens S, Shamsi S. 2020. Morphological and genetic characterisation of the introduced copepod *Lernaea cyprinacea* Linnaeus (Cyclopoida: Lernaeidae) occurring in the Murrumbidgee catchment, Australia. Mar. Freshw. Res. 72: 876–886.

