## Supplementary material for "Host-parasite interactions in perpetual darkness: macroparasite diversity in the cavefish *Astyanax mexicanus*": Table S1

| **Table S1**. Primers used for PCR and sequencing of parasites in this study. | | | | |
| --- | --- | --- | --- | --- |
| Genetic region | Primer name | F/ R | Sequence (5'− 3') | Reference |
| cox1 | JB3/ Asmit1 | F | TTTTTTGGGCATCCTGAGGTTTAT | Bowles et al. 1992 |
|  | JB4.5 | R | TAAAGAAAGAACATAATGAAAATG | Bowles et al. 1992 |
|  | Schisto3 | R | TAATGCATMGGAAAAAAACA | Lockyer et al., 2003 |
|  | 507 | F | AGTTCTAATCATAARGATATYGG | Nadler et al., 2006 |
|  | HCO2198 | R | TAAACTTCAGGGTGACCAAAAAATCA | Folmer et al., 1994 |
|  | pr-b | R | AGTTCTAATCATAARGATATYGG | Bessho et al., 1992 |
| cox2 | 210 | F | TTTTCTAGTTATATAGATTGRTTYAT | Nadler & Hudspeth, 2000 |
|  | 211 | R | CACCAACTCTTAAAATTATC | Nadler & Hudspeth, 2000 |
| 28S | 391 | F | AGCGGAGGAAAAGAAACTAA | Nadler & Hudspeth, 1998 |
|  | 536 | R | CAGCTATCCTGAGGGAAAC | García-Varela & Nadler, 2005 |
|  | 502* | F | CAAGTACCGTGAGGGAAAGTTGC | García-Varela & Nadler, 2005 |
|  | 503 | R | CCTTGGTCCGTGTTTCAAGACG | Nadler et al., 2003 |
|  | U178 | F | GCACCCGCTGAAYTTAAG | Lockyer et al., 2003 |
|  | 1200R* | R | GCATAGTTCACCATCTTTCGG | Lockyer et al., 2003 |
|  | L1642 | R | CCAGCGCCATCCATTTTCA | Lockyer et al., 2003 |
| F: forward primer; R: reverse primer | | | | |
| Asterisk indicates primers used only in sequencing reaction. | | | | |

**REFERENCES**

Bessho Y, Ohama T, Osawa S. 1992. Planarian mitochondria I. Heterogeneity of cytochrome c oxidase subunit I gene sequences in the	freshwater planarian, *Dugesia japonica*. *J. Mol. Evol*. **34**: 324–330.

Bowles J, Blair D, Mcmanus D. 1992. Genetic variants within the genus *Echinococcus* identified by mitochondrial DNA sequencing. *Mol.	Biochem. Parasitol.* **54**:165–173.

Folmer O, Black M, Hoeh W, Lutz R, Vrijenhoek R. 1994. DNA primers for amplification of mitochondrial cytochrome c oxidase subunit I from	diverse metazoan invertebrates. *Mol. Marine Biol. Biotechnol*. **3**: 294–299.

García-Varela M, Nadler SA. 2005. Phylogenetic relationships of palaeacanthocephala (Acanthocephala) inferred from SSU and LSU rDNA	gene sequences. *J. Parasitol*. **91**: 1401–1409.

Lockyer AE, Olson PD, Littlewood DTJ. 2003. Utility of complete large and small subunit rRNA genes in resolving the phylogeny of the	Neodermata (Platyhelminthes): Implications and a review of the cercomer theory. *Biol. J. Linn. Soc*. **78**: 155–171.

Nadler SA, Bolotin E, Stock SP. 2006. Phylogenetic relationships of *Steinernema* Travassos, 1927 (Nematoda: Cephalobina: Steinernematidae)	based on nuclear, mitochondrial and morphological data. *Syst. Parasitol*. **63**: 161–181.

Nadler A, Carreno RA, Adams BJ, Kinde H, Baldwin JG, Mundo-Ocampo M. 2003. Molecular phylogenetics and diagnosis of soil and clinical isolates of *Halicephalobus gingivalis* (Nematoda: Cephalobina: Panagrolaimoidea), an opportunistic pathogen of horses. *Int. J. Parasitol*. **10:** 1115–1125.

Nadler SA, Hudspeth DSS. 1998. Ribosomal DNA and Phylogeny of the Ascaridoidea (Nemata: Secernentea): Implications for Morphological	Evolution and Classification. *Mol. Phylogenet. Evol*. **10**: 221–236.
