## Supplementary material for "Host-parasite interactions in perpetual darkness: macroparasite diversity in the cavefish *Astyanax mexicanus*": Table S2

| **Table S2.** Primer combination and PCR conditions. | | | | |  |
| --- | --- | --- | --- | --- | --- |
| Genetic marker | PCR + SEQ | SEQ | Tm (ºC) | Parasite group | Size of product (pb) |
| cox1 | JB3− JB4.5 | − | 42 | N; T; M | ~ 400 |
|  | 507− HCO | − | 48 | N | ~ 630 |
|  | 507− prb | − | 48 | N | ~ 1014 |
|  | Asmit− Schisto3 | − | 50 | M | ~ 480 |
|  | pra− prb | − | 40 | N | ~ 400 |
| cox2 | 210− 211 | − | 46 | N | ~ 550 |
| 28S | 391− 536 | 502, 503 | 52 | N; T | ~ 1149 |
|  | U178− L1642 | 1200R | 52 | M | ~ 1389 |
| 18S | G18S4− 649 | 136 | 54 | N | ~ 1600 |
| T; trematoda; M: monogenea; N: nematoda | | | | | |
